# Identifying conservation priority areas for the most under-protected globally threatened birds in Latin America

**DOI:** 10.1101/2025.05.09.653075

**Authors:** Marcelo F. Tognelli, David A. Wiedenfeld, Daniel J. Lebbin, Amy Upgren, Michael J. Parr

**Affiliations:** American Bird Conservancy, PO Box 249, The Plains, VA 20198 USA

**Keywords:** Gap analysis, area of habitat, minimum conservation target, prioritization, Latin America

## Abstract

In this study, we assess how well globally threatened bird species in Latin America are currently represented within existing protected areas, and we identify conservation priority areas for the most under-protected species. We used publicly available data to map the Area of Habitat (AOH) for each of the 149 land birds in South America listed on the IUCN Red List of Threatened Species as either Critically Endangered, Endangered, or Vulnerable (the latter only under criterion D – those with the smallest populations or ranges). We then set two minimum conservation targets for each species: first a population-based target that estimates the proportion of the AOH needed to conserve 1,000 mature individuals (or the total population if smaller) to prevent extinction; and second, an area-based target that estimates the amount of habitat needed to sustain the long-term conservation of each species. The AOH maps were then overlaid with existing protected areas to identify those species that do not yet have their minimum conservation targets met. Using this approach, we identified 10 species that require additional protection to avoid extinction, and a further 54 species that need expanded protection for sustained conservation. Fortunately, the majority of species already met their target. We also ran a prioritization analysis to identify and map the places most important for meeting the goals of the under-protected species. We found that just 661.4 km^2^ is needed to meet population-based targets for the ten species of greatest concern, and 16,360 km^2^ is needed to meet area-based targets for all under-protected species combined. These areas represent <0.1% of the region’s land area and are mostly concentrated in five countries (Mexico, Colombia, Ecuador, Peru, and Brazil). Expanding reserves to cover these areas should result in both improved conservation outcomes and Red List status for these species.

## Introduction

Recent global assessments have analyzed and synthesized data showing the staggering loss of biodiversity and ecosystem deterioration due to human actions (Díaz et al. 2019, Secretariat of the Convention on Biological Diversity 2020, WWF 2022). Latin America has shown the greatest regional decline in mean vertebrate population abundance, at 94% between 1970 and 2018 (WWF 2022). The majority of globally threatened species are declining due to land use change (Díaz et al. 2019; Harfoot et al. 2021; Jaureguiberry et al. 2022), and protecting terrestrial biodiversity and preventing extinctions therefore requires protection and management of habitat. Numerous conservation organizations and global initiatives are working towards this goal. Target 3 of the Kunming-Montreal Global Biodiversity Framework (KMGBF) aims to “Ensure and enable that by 2030 at least 30 per cent of terrestrial, inland water, and of coastal and marine areas, especially areas of particular importance for biodiversity and ecosystem functions and services, are effectively conserved and managed,” commonly known as 30×30 initiative (Kunming-Montreal Global Biodiversity Framework; CBD 2022), and many governments and private organizations are working towards fulfilling this target, as well as other national and local goals for habitat protection.

Identifying biodiversity conservation priority areas to prevent species’ extinction is an important step needed to inform strategic investment in conservation. Key Biodiversity Areas (KBAs, sites of significance for the global persistence of biodiversity; IUCN 2016), including Alliance for Zero Extinction sites (AZE, sites supporting the overwhelming majority of the remaining population of one or more Critically Endangered or Endangered species; Alliance for Zero Extinction 2013,) have been identified for this purpose. Efforts are underway to update and increase the comprehensiveness of these site networks (KBA Partnership 2023). Conserving key sites is a critical tool to help deliver Target 4 of the KMGBF on undertaking urgent management actions to prevent species extinctions (Boyd et al. 2008; Bolam et al. 2023). By understanding the distribution of specific species and their overlap with existing protected areas, conservation practitioners can pinpoint which highly threatened species are under-protected and where to invest to fill the gaps in the existing protected area network to prevent species extinctions.

Birds are the best known of any class of vertebrates, and previous studies have overlaid the distributions of bird species with protected area data to determine the proportion of each species’ range that is already protected (Venter et al. 2014; Butchart et al. 2015). However, these global assessments have used the species’ mapped ranges, rather than the area of habitat (AOH) within them, which provides a finer-resolution indication of species’ potential distributions (Beresford et al. 2011, Brooks et al. 2019). The main sources of range maps for this purpose are the International Union for Conservation of Nature Red List of Threatened Species’ (the “Red List”) distribution maps, produced by BirdLife International in its role as the IUCN Red List Authority for birds (hereafter Red List range maps). Recent efforts to improve these maps use high-resolution remote sensing and citizen science data (e.g., Huang et al. 2021, Palacio et al. 2021). Such citizen science data allow the refinement of bird distribution maps, including the creation of AOH maps using various data sources and methods (Beresford et al. 2011; Ocampo-Peñuela et al. 2016, Huang et al. 2021, Palacio et al. 2021; Lumbierres et al. 2022; Warudkar et al. 2022; Medina et al. 2024).

Some recent range-mapping efforts have relied on the use of occurrence data (i.e., point localities from museum specimen records and/or citizen science projects such as eBird, iNaturalist, etc., often through the Global Biodiversity Information Facility data infrastructure network) to update or improve the existing Red List range maps (Huang et al. 2021; Warudkar et al. 2022) or to derive new ranges based on species’ extent of occurrence (EOO) (Palacio et al. 2021). However, most of these efforts have focused only on subsets of species, such as forest birds. We assessed these approaches to standardize the process across all species and habitats and determine the most appropriate approach for threatened birds of the Americas.

Given the issues and lessons learned from these efforts, we aimed to systematize the generation of AOH maps with minimal introduction of errors. Our approach includes some of the features of the other methods, tempered with expert opinion. Here we use publicly available bird occurrence data to develop a method to map the Area of Habitat (AOH), of 149 terrestrial land birds in Latin America, and overlay these with existing protected areas as compiled by the World Database of Protected Areas (WDPA, UNEP-WCMC and IUCN 2023), plus additional private protected areas not gazetted. We aim to quantify how much habitat is protected and unprotected, set habitat protection goals for under-protected species, identify which species are under-protected, and map areas where additional habitat protection projects could be developed and implemented to fill in the gaps of existing protected areas and reduce the extinction risk for these species.

## Methods

### Species data

We focused on species of land birds from Latin America (excluding the Caribbean and other outlying islands) listed in the Red List categories “Critically Endangered” (CR), “Endangered” (EN), and “Vulnerable” (VU D1 and D2) (IUCN 2022) as of June 2024. We included only VU species listed under Red List criterion D (i.e., D1, and D2; see IUCN 2012 for definition of criteria) because these are species that have populations of < 1,000 mature individuals and/or have very restricted distributions and can become CR or Extinct in a very short period of time. We also excluded seven species that are considered “lost” (i.e., they have not been reported in at least ten years; https://searchforlostbirds.org/), and Great-billed Seed-finch *Sporophila maximiliani*, because its taxonomy is currently so unclear that its occurrence and distribution cannot be properly determined. After applying these filters, the final set included 149 threatened bird species.

We obtained point locality data for each of the 149 selected bird species from eBird (eBird Basic Dataset. Version: EBD_relDec-2022. Cornell Lab of Ornithology, Ithaca, New York. Dec 2022, accessed: February 14, 2023). We filtered the eBird data to occurrences from the year 2000 onward to have only recent data and likely extant populations; only those using the “Traveling” Protocol Type with a distance covered by the observer of < 5 km; and those with the Protocol Type of “Stationary” and “Incidental.” This limits the data to recent records likely to have the most precise location information.

### Habitat, elevation, land conversion pressure, and protected area data

We obtained land cover data from the European Space Agency – Climate Change Initiative (ESA-CCI) Land Cover map for the year 2020 (v. 2.1.1.; http://maps.elie.ucl.ac.be/CCI/viewer/index.php; accessed on May 6, 2023). The layer consists of 22 primary classes and 15 additional sub-classes of land cover at 300 m spatial resolution. Moreover, given the challenge that represents accurately mapping and differentiating natural and semi-natural grasslands versus cultivated areas, we used two recently developed layers to account for areas that are croplands: the Global Cropland-Extent Product (GCEP30, Thenkabail et al. 2021) and the Global Grassland Class and Extent maps (Parente et al. 2024). The native resolution of both these layers is 30 m, so they were resampled to 300 m using nearest neighbor interpolation to use the same resolution across all layers.

We used elevation data from the Shuttle Radar Topography Mission (SRTM 4.1) digital elevation model layer (downloaded from https://srtm.csi.cgiar.org/srtmdata/; accessed on June 16, 2023; 250 m resolution). The elevation layer was resampled to 300 m using bilinear interpolation to use the same spatial resolution as the habitat layer.

To account for local habitat degradation in our prioritization analysis, we used the Conversion Pressure Index (CPI) layer developed by Oakleaf et al. (2024). The CPI is a global map of land conversion that combines past rates of anthropogenic pressure (derived from multiple drivers) with suitability maps for potential future expansion by large-scale development projected up to 2030 (Oakleaf et al. 2024). CPI values range from 0, which indicates no conversion pressure, to 1, which represents the highest conversion pressure. According to the authors, this information can be used to identify areas for proactive conservation to ensure national commitments under the KMGBF. The CPI layer was resampled to 300 m using bilinear interpolation to use the same spatial resolution as the other layers.

We downloaded protected area data from the World Database of Protected Areas (WDPA; UNEP-WCMC and IUCN 2024; www.protectedplanet.org; accessed on April 9, 2024). The WDPA includes different protected area management categories developed by the IUCN, but we included all categories in our analysis as the reporting of categories and their enforcement is widely variable among countries. We also added several private protected areas and/or protected area expansions that are owned and managed by American Bird Conservancy (ABC) partners, but which are not yet recognized in the WDPA because they have been established recently. We merged the overlapping protected area polygons into a single polygon to avoid overestimation of the protected area coverage. We did not include Other Effective area-based Conservation Measures (OECMs; CBD 2018) as few Latin American countries have recognized any sites as OECMs yet. Colombia is the only country to have done so to date, and the effectiveness of these sites has not yet been assessed.

### Developing Area of Habitat (AOH) maps

The method for estimating AOH involves selecting the available suitable habitat(s) and elevation range within the species’ mapped geographic range (Brooks et al. 2019). The geographic range is defined as the “current known limits of distribution of a species, accounting for all known, inferred or projected sites of occurrence” (IUCN 2016) and is depicted as a polygon map in the IUCN Red List (Red List range maps). These maps are derived from a variety of sources, including specimen localities obtained from museum data, observer records documented in Red Data Books, published literature, survey reports and other unpublished sources; records of species documented in Important Bird and Biodiversity Area assessments, distribution atlases derived from systematic surveys, distribution maps in field guides and other handbooks, and expert opinion (Buchanan et al 2011).

While eBird and other citizen science data have been used in some cases, this has not happened systematically for all species. Some of the range maps are outdated, as they are typically updated only when the species’ risk of extinction is re-assessed. For instance, almost half of the species included in our analysis had Red List assessments that were at least five years old, and therefore likely have range maps at least five years old. Given this, we mapped the existing Red List range maps together with the eBird locality data, and used these inputs to modify the ranges (whenever needed) based on expert opinion (see Figure 1 for an example).

**Figure 1.**
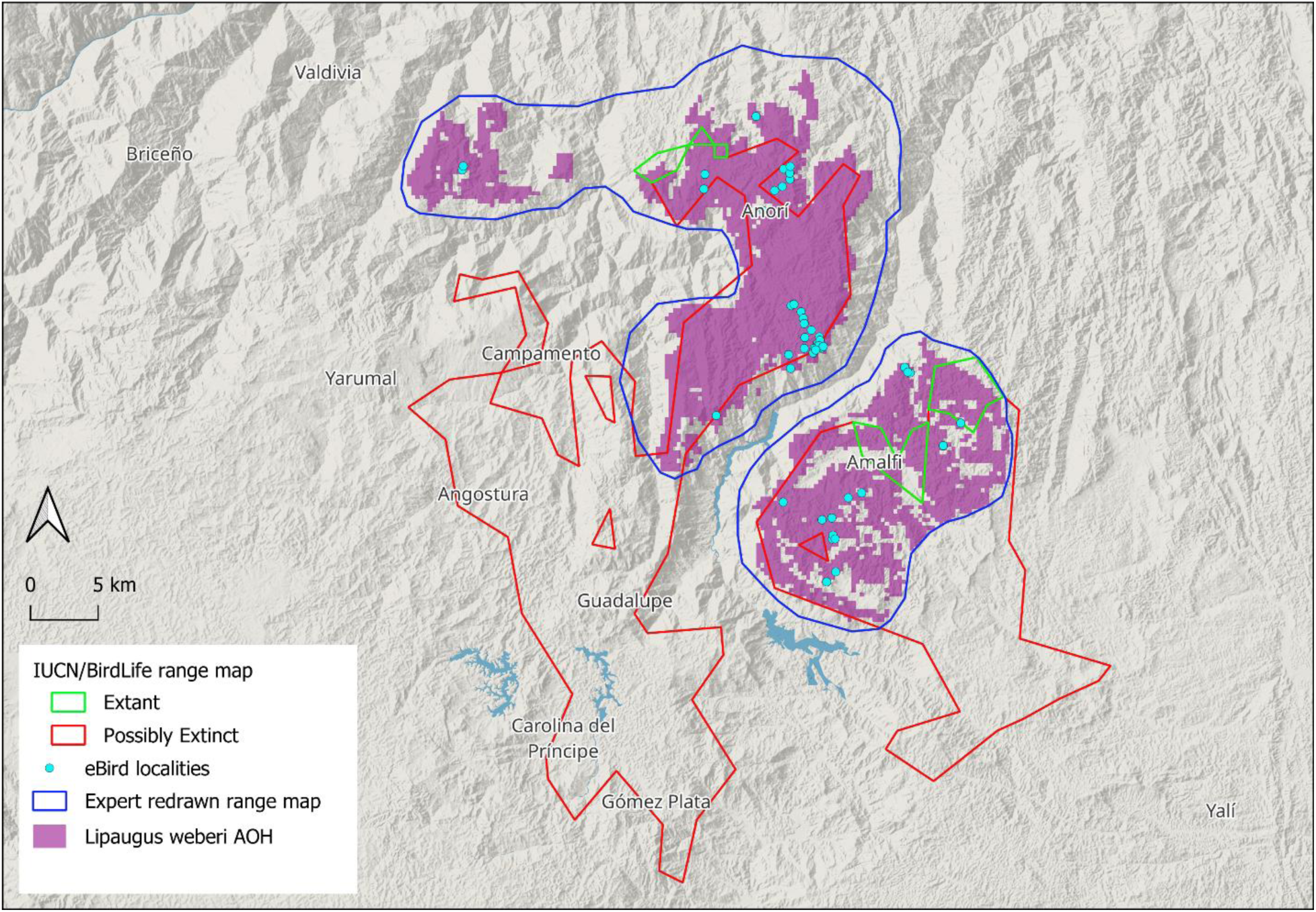
Example map, for Chestnut-capped Piha (*Lipaugus weberi*) in western Colombia. The map shows the eBird localities (turquoise dots), the original IUCN/BirdLife range map (showing the Extant and Possibly Extinct portions of the range as green and red polygons respectively), the new range created in this study (blue polygons), and the AOH (purple) generated within the new range map by selecting the species’ suitable habitat and elevation range.

To determine the elevation range of the species, we used the range of elevations from the existing eBird data points, excluding the highest and lowest 1% to reduce the impact of outliers. Once the new ranges were delimited, we overlaid the point locality data with the habitat (land cover) layer to extract the corresponding values for each point. We checked the habitat classes against the habitat coding and descriptions in the IUCN Red List assessments and specific publications, and excluded those that were not listed or mentioned there. Also, because the land cover map may misclassify some pixels, the land cover classes for all presence data were inspected visually and those that were obvious mistakes were excluded (a common mistake in satellite image classification is classifying pastures or certain crops as natural grasslands). Based on this information, we proceeded to derive AOH maps by intersecting the elevation and habitat layers within the new range map, and removing the pixels of unsuitable habitats (including the cropland layers) and outside the elevation range. All analyses and maps were done with QGIS (ver. 3.34; www.qgis.org).

### Minimum Conservation Targets and Gap Analysis

To guide future avian conservation priorities in the Americas, we defined two minimum conservation targets (i.e., the minimum proportion of a species’ AOH that should be included in protected areas to be considered adequately represented and safeguarded) for protected area coverage for each focal species. The first area-based target estimated the proportion of the AOH needed to be represented in protected areas in order to sustain conservation of those species. The second population-based target estimated the area needed to protect 1,000 mature individuals of each species (or the maximum population size for those species whose estimated maximum population is < 1,000 mature individuals) in order to prevent imminent extinction of these species.

The minimum conservation target based on the proportion of AOH was set to 100% for species with AOH ≤ 50 km^2^, whereas for species with AOH > 20,000 km^2^, the minimum conservation target was set to 4% of the range. These values and cutoffs were selected to ensure that the entire population is protected for species with small ranges and to ensure an area adequate for conservation to protect for species with large ranges. For all other species (in the range of >50 km^2^ and <= 20,000 km^2^), the target was interpolated using a linear regression on the log-transformed AOH (Rodrigues et al. 2004).

The population-based target to protect was selected using the population size threshold for a species to avoid qualifying for the Vulnerable category under criterion D1 of the IUCN Red List of Threatened Species (IUCN 2012), which is 1,000 mature individuals. To calculate the area needed to include 1,000 mature individuals, we estimated the territory sizes or area requirements for each species in our list as described in Table 1. Estimates were obtained from the literature (see Supplementary Information). Whenever there was no direct information for the species in our list, we used data from congeneric species of similar size and habits.

**Table 1.**
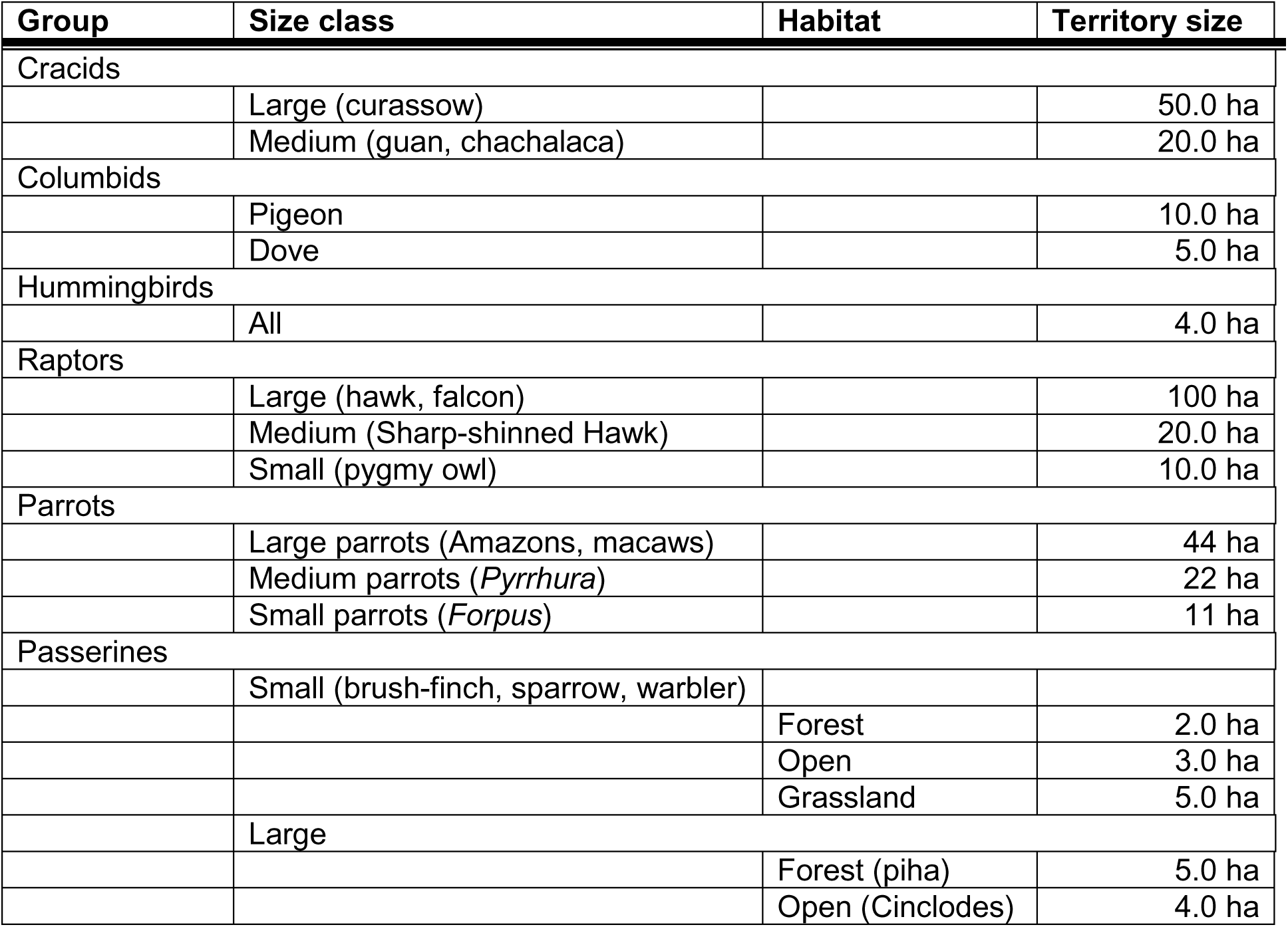
Territory size estimates for birds, by size class. Estimates are per individual; therefore, the area must be doubled for a pair, or multiplied by average flock size for species such as parrots that travel in flocks.

| Group | Size class | Habitat | Territory size |
| --- | --- | --- | --- |
| Cracids |  |  |  |
|  | Large (curassow) |  | 50.0 ha |
|  | Medium (guan, chachalaca) |  | 20.0 ha |
| Columbids |  |  |  |
|  | Pigeon |  | 10.0 ha |
|  | Dove |  | 5.0 ha |
| Hummingbirds |  |  |  |
|  | All |  | 4.0 ha |
| Raptors |  |  |  |
|  | Large (hawk, falcon) |  | 100 ha |
|  | Medium (Sharp-shinned Hawk) |  | 20.0 ha |
|  | Small (pygmy owl) |  | 10.0 ha |
| Parrots |  |  |  |
|  | Large parrots (Amazons, macaws) |  | 44 ha |
|  | Medium parrots (*Pyrrhura*) |  | 22 ha |
|  | Small parrots (*Forpus*) |  | 11 ha |
| Passerines |  |  |  |
|  | Small (brush-finch, sparrow, warbler) |  |  |
|  |  | Forest | 2.0 ha |
|  |  | Open | 3.0 ha |
|  |  | Grassland | 5.0 ha |
|  | Large |  |  |
|  |  | Forest (piha) | 5.0 ha |
|  |  | Open (Cinclodes) | 4.0 ha |

To assess the level of protection of each species, we intersected the individual AOH maps with the World Database of Protected Areas layer and estimated the area and proportion within protected areas. We were then able to assess the conservation gap (measured in squared kilometers) for each focal species and considering the two conservation targets for each species.

To find the main priority areas for conservation, we ran the Zonation 5 v2.2 software (Moilanen et al. 2022). Zonation is a spatial prioritization software for conservation planning that identifies which locations in a landscape are more important for retaining biodiversity. The advantage of Zonation (versus other prioritization programs) is that it can be used with high-resolution continental analysis on a modest computer system. The main output is a priority rank map where grid cells are ranked from highest to lowest priority in terms of conservation value. Therefore, the user can select the top X% ranked cells of the entire study area, depending on the objective of the prioritization. Because the new version of Zonation does not include a target-based planning algorithm, instead of selecting a predetermined percentage of ranked cells for all species, we selected the percentage of area represented for each individual species, using their minimum conservation targets as a cut-off criterion. For instance, if the minimum conservation target of a particular species is 27% of its AOH, we selected the top X% ranked cells that included 27% of that species’ AOH, and so forth for all priority species. We used the protected areas layer as a hierarchical mask to lock in those cells in the solution and identify the priority areas outside the present protected area network that will be needed to reach the species’ minimum conservation targets. We also used the Conversion Pressure Index (described above) as a condition layer (Moilanen et al. 2024), that accounts for local habitat deterioration and its influence on the prioritization process. The purpose here is to identify preliminary areas for conservation at a large scale. Actual implementation of habitat protection should refine this analysis with more detailed information at the landscape scale (including existing OECMs).

## Results

We redrew the range maps and mapped the Area of Habitat for the 149 priority threatened bird species in Latin America. See Figure 1 depicting an example map for the Chestnut-capped Piha (*Lipaugus weberi*), endemic to northwestern Colombia. Of the 149 species included in our analysis, 20 are in the IUCN Red List Category Vulnerable (VU D1 or D2), 94 Endangered (EN), and 35 Critically Endangered (CR).

When assessing the minimum conservation targets based on the proportion of AOH to be protected, over half of the species (85) have their targets met (i.e., 100% of the proposed minimum conservation area is currently protected), and 64 need to increase the area under protection to reach their targets (Table S1). On average, 37.9% of the AOH of all species analyzed is currently under protection, ranging from four species with less than 1% protected (Little Inca-finch *Incaspiza watkinsi*, Chilean Woodstar *Eulidia yarrelli*, Oaxaca Hummingbird *Eupherusa cyanophrys*, and Marañon Antshrike *Thamnophilus shumbae*) to nine with > 99% of their AOH protected (Table S1). The average conservation gap for the 64 species that do not have their proportion of AOH target protected is 12.1%.

The main priority areas for covering the conservation gaps of the 64 species whose targets have not been met are in the Tropical Andes of Peru, Ecuador, and Colombia; northeastern Brazil; the Caribbean coast of Colombia; and central and northern Mexico (Figure 2). The area that would be needed to meet the minimum conservation targets for these species, without considering the overlap in multiple species distributions, is 22,406 km^2^. However, in some cases, multiple species may occur in the same area; therefore, protecting a specific area may protect more than one species. Because the Zonation algorithm produces priority solutions that are well-balanced and where priority sites have high complementarity (Moilanen et al. 2022), the total area needed to meet the minimum conservation target for the 64 focal species is reduced to 16,360 km^2^.

**Figure 2.**
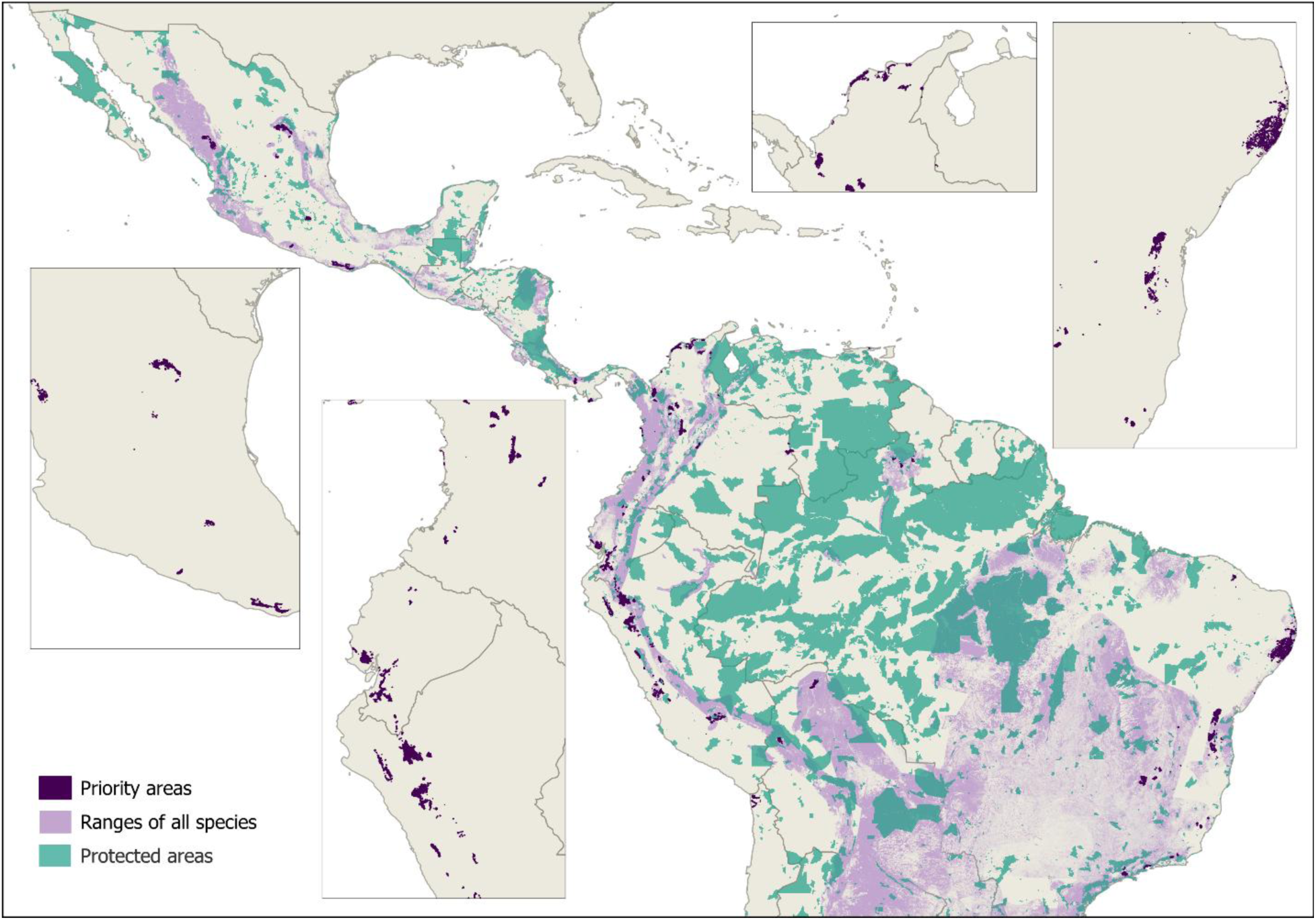
The existing protected area network (green), the range maps of all focal species (light lilac), and the priority areas (dark purple) needed to meet the proposed minimum conservation targets for all 64 species for which the minimum conservation targets are not already met. Protected areas are excluded in the enlarged inset maps for better visualization.

An example of the Zonation software results identifying areas with overlap of several species can be seen in the Murici area, in the State of Alagoas, northeastern Brazil (Figure 3), where a conservation priority area of 5,234 ha contains parts of the AOH of ten of the species comprised in this study, including nine that do not have their minimum conservation targets met (Forbes’s Blackbird *Anumara forbesi*, Pernambuco Foliage-gleaner *Automolus lammi*, White-collared Kite *Leptodon forbesi*, Scalloped Antbird *Myrmoderus ruficauda*, Alagoas Antwren *Myrmotherula snow*i, Alagoas Tyrannulet *Phylloscartes ceciliae*, Pinto’s Spinetail *Synallaxis infuscata*, Orange-bellied Antwren *Terenura sicki*, and Long-tailed Woodnymph *Thalurania watertonii*). This relatively small area would protect an average of 11.1% (range 0.5 – 54.3 %) of the AOH of these nine species.

**Figure 3.**
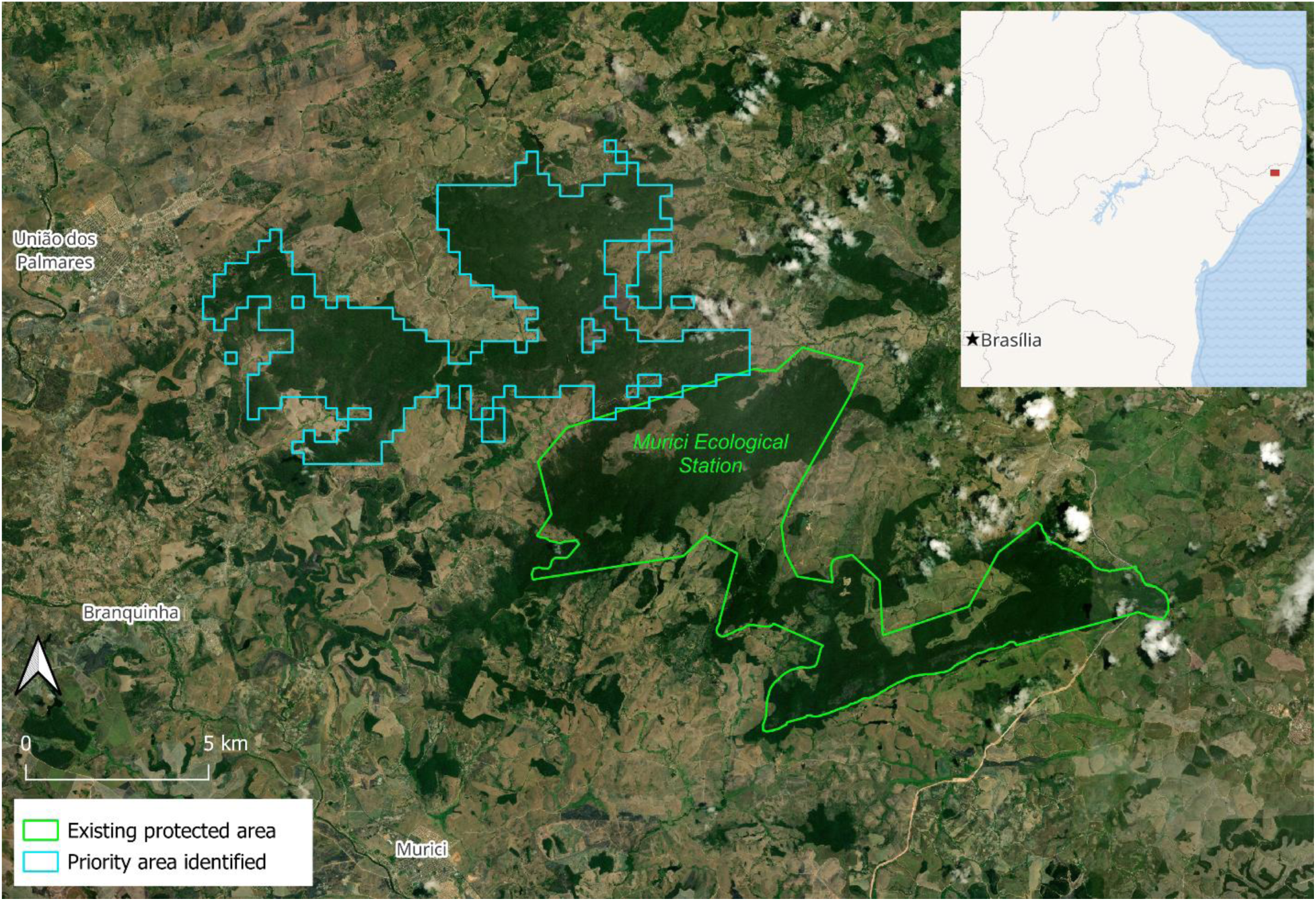
Priority conservation area of 5,234 ha identified by the Zonation software in the Murici area, northeastern Brazil (inset map). This area would protect an average of 11.1% of the AOH of nine priority bird species.

When assessing the minimum conservation targets based on the area needed to protect up to 1,000 mature individuals of each species, 139 (93%) of the species have their targets met, and 10 need to increase the area under protection to reach their targets (Table S1). The average conservation gap for these 10 species is 8.3% (range 0.3% - 36.3%), and the total area needed to cover these gaps is 661.4 km^2^ (range 0.9 km^2^ - 236.9 km^2^). Because these 10 species are a subset of the 64 species identified using the proportion of AOH conservation target, the priority areas for them can be selected using the broader scale prioritization exercise.

## Discussion

Our results identified 64 under-protected bird species that require 16,360 km^2^ of additional habitat protection within very discrete areas to meet their minimum conservation targets. This is a tiny fraction (>0.1%) of the total area of approximately 20,000,000 km^2^ of Central and South America. The distribution of the lands needing protection is not even across the Neotropics, and is concentrated mostly in five countries (Mexico, Colombia, Ecuador, Peru, and Brazil), but even within these countries, the area is not a significant fraction of their total area. For example, it represents approximately 0.14% of the total area of Mexico, 0.24% of Colombia, 0.74 % of Ecuador, 0.34% of Peru, and 0.04% of Brazil.

An important feature of our analysis is that it illustrates protection gaps for globally threatened birds, providing useful information to ensure minimum protection for and prevent extinctions of this important subset of species. The 16,360 km^2^ identified represents less than 0.1% of the region’s 30×30 protection goals. Increased public commitment to habitat protection, exemplified by national commitments to the 30×30 Initiative, are providing the enabling conditions to allow these lands to become conserved. Therefore, we recommend that the areas identified in this paper are prioritized for conservation within the 30×30 Initiative, as well as included in countries’ biodiversity-inclusive spatial plans (Target 1 of the KMGBF), and be considered in plans for restoration under Target 2. They can also be included in countries’ National Biodiversity Strategies and Action Plans (NBSAPs), and other biodiversity conservation efforts. Further implementation of habitat protection within these priority areas to meet the conservation thresholds for individual species will require additional planning and project development at a local landscape scale with additional land tenure information.

As expected, the priority sites identified in our study overlap with sites identified as internationally significant for biodiversity. For instance, 35.6% of the 16,360 km^2^ falls within the global network of KBAs, overlapping with 108 KBAs and covering an average 14.5% (range 0.1 - 78.9%) of the area of each of these 108 sites. Most importantly, besides including 665 bird KBA-qualifying species (i.e. species for which the site meets KBA criteria), these sites also include 305 KBA-qualifying species from other taxonomic groups, such as 125 amphibians, 83 plants, 50 reptiles, 20 mammals, 13 freshwater fish, and 14 arthropods. Moreover, 29 of the 108 KBAs are AZE sites that harbor the majority of the remaining population of one or more Critically Endangered or Endangered species, including 38 non-bird species.

Although we consider 86 of the 149 species to meet their minimum proportional area conservation target, conservation efforts are still necessary to ensure that the protected areas are appropriately managed so that habitats and species are effectively protected within these existing protected areas.

The recent analysis for the Northern Andes by Medina et al. (2024) is complementary to our work. Our analysis focuses on species already considered threatened from all of Latin America outside the Caribbean, whereas their analysis includes all range-restricted (< 50,000 km^2^) species and focuses on a relatively small, but important area. Medina et al. (2024) use a method similar to that of Huang et al. (2021), with its reliance on the original IUCN/BirdLife maps and treatment of absence data. Although our two analyses differ, they are nonetheless similar in important ways, identifying some of the same regions as key areas. This is a significant result, indicating that these areas are important in a fundamental biological way, at least in the Northern Andes region, an area that has been highlighted as a conservation priority for threatened resident species of mammals, birds, reptiles, and amphibians (Wilson et al. 2021).

The 149 bird species in our analysis represent 13 orders of birds, with diverse life histories and ecologies, occupying landscapes ranging from sea level and relatively unbroken lands to high-elevation and highly dissected mountainous regions. They occur in habitats ranging from very humid forests to semi-arid deciduous woodlands. Developing methods to produce range maps that can account for the myriad differences among species and conditions has proven difficult, as seen from the fact that several new methods for producing range maps for birds have been proposed in just the last three years. An additional challenge for producing maps for many species in the Neotropics, especially ones that are threatened and therefore already rare, is the lack of data and the biased distribution of data that do exist. Citizen science programs such as eBird and scientific data collection efforts such as GBIF are very important sources of bird distribution information but often do not provide sufficient data from across a representative portion of a species’ range to develop highly accurate range maps. Therefore, expert opinion remains important in the production of bird range maps.

## Supporting information

References Territory Size

## Acknowledgments

We kindly thank David Luther, Stuart Butchart, Ryan Huang, Wilderson Medina, James Lewis, and Jorge Velásquez-Tibatá for providing feedback on early versions of this manuscript. Our thinking on this paper also benefitted from numerous discussions with Stuart Pimm, Ryan Huang, Wil Medina, and others in their working group over the last several years.

## Supplementary Materials

*Table S1*. List of species included in the analysis, including their common names, threatened category, AOH area, level of protection, minimum conservation targets, additional AOH needed to reach the minimum conservation targets (both in % and km^2^), and percentage of a species’ AOH over the minimum conservation target that was included in the Zonation solution. *Supplementary information*. List of references that was used for territory size information.

