## Supplementary material for "Identifying conservation priority areas for the most under-protected globally threatened birds in Latin America": References Territory Size

Supplementary Information

BirdLife International. 2018. Grallaria fenwickorum. The IUCN Red List of Threatened Species 2018: e.T22736345A131861216. Accessed on 29 April 2025.

BirdLife International. 2021. *Merulaxis stresemanni*. The IUCN Red List of Threatened Species 2021: e.T22703477A161689029. Accessed on 29 April 2025.

Dahlin, C. R., Blake, C., Rising, J. and Wright, T. F. 2018. Long-term monitoring of Yellow-naped Amazons (*Amazona auropalliata*) in Costa Rica: breeding biology, duetting, and the negative impact of poaching. Journal of Field Ornithology 89: 1–10.

Dolkas, G. A. and Neiman, T. J. 2020. Tufted Tit-Tyrant (*Anairetes parulus*), version 1.0. Birds of the World. https://doi.org/10.2173/bow.tuttyr1.01

Gill, F. B. 1988. Trapline Foraging by Hermit Hummingbirds: Competition for an Undefended, Renewable Resource. Ecology 69: 1933–1942.

Greeney, H. F. 2020. Variegated Antpitta (*Grallaria varia*), version 1.0. Birds of the World. https://doi.org/10.2173/bow.varant2.01

Gussoni, C. O., Fitzpatrick, J. W., Sharpe, C. J. and Spencer, A. J. 2021. Alagoas Tyrannulet (*Phylloscartes ceciliae*), version 2.0. Birds of the World. https://doi.org/10.2173/bow.alatyr1.02

Haggerty, T. M. and Morton, E. S. 2020. Carolina Wren (*Thryothorus ludovicianus*), version 1.0. Birds of the World. https://doi.org/ 10.2173/bow.carwre.01

Johnson, L. S. 2024. Northern House Wren (*Troglodytes aedon*), version 1.1. Birds of the World. https://doi.org/ 10.2173/bow.houwre.01.1

Krabbe, N. and Juiña, M. 2020. Pale-headed Brushfinch (*Atlapetes pallidiceps*), version 1.0. Birds of the World. https://doi.org/ 10.2173/bow.phbfin1.01

Mueller, A. J. 2020. Inca Dove (*Columbina inca*), version 1.0. Birds of the World. https://doi.org/ 10.2173/bow.incdov.01

Pavan, L. I., Jankowski, J. E. and Hazlehurst, J. A. 2020. Patterns of territorial space use by Shining Sunbeams (*Aglaeactis cupripennis*), tropical montane hummingbirds. Journal of Field Ornithology 91: 1–12.

Potter, A. B. 2020. Gray-winged Trumpeter (*Psophia crepitans*), version 1.0. Birds of the World. https://doi.org/ 10.2173/bow.gywtru1.01

Remsen, J. and Sharpe, C. J. 2020. Royal Cinclodes (*Cinclodes aricomae*), version 1.0. Birds of the World. https://doi.org/ 10.2173/bow.roycin1.01

Rivas-Fuenzalida, T., Grande, J. M., Kohn, S., Vargas, F. H., and Zuluaga Castañeda, S. 2024. Black-and-chestnut Eagle (*Spizaetus isidori*), version 3.0. In Birds of the World (S. M. Billerman, Editor). Cornell Lab of Ornithology, Ithaca, NY, USA. https://doi.org/10.2173/bow.baceag2.03

Rueda-Uribe, C., Sargent, A. J., Echeverry-Galvis, M. Á., Camargo-Martínez, P. A., Capellini, I., Lancaster, L. T., Rico-Guevara, A. and Travis, J. M. J. 2024. Tracking Small Animals in Complex Landscapes: A Comparison of Localisation Workflows for Automated Radio Telemetry Systems. Ecology and Evolution 14: e70405.

Schulenberg, T. S. 2020. Amazonian Pygmy-Owl (*Glaucidium hardyi*), version 1.0. In Birds of the World (T. S. Schulenberg, Editor). Cornell Lab of Ornithology, Ithaca, NY, USA. https://doi.org/10.2173/bow.amapyo1.01

Small, M. F., Fuller, J. C., Hook, M. W. and Dukes, W. F. 2021. Does Amount of Urban Area Around Predominantly Rural Banding Sites for Mourning Doves Affect Harvest in the Carolinas? Journal of the Southeastern Association of Fish and Wildlife Agencies 8: 84–88.

Stahala, C. 2008. Seasonal movements of the Bahama Parrot (*Amazona leucocephala bahamensis*) between pine and hardwood forests: implications for habitat conservation. The Neotropical Ornithological Society 19: 165–171.

Suzuki, I., Fearnside, N., Tori, W. and Pareja, J. I. 2020. Screaming Piha (*Lipaugus vociferans*), version 1.0. Birds of the World. https://doi.org/ 10.2173/bow.scrpih1.01

Udoye, K. C. and Schulenberg, T. S. 2020. Razor-billed Curassow (*Mitu tuberosum*), version 1.0. Birds of the World. https://doi.org/ 10.2173/bow.rabcur2.01

Zimmer, K., Isler, M. L. and Kirwan, G. M. 2020. Ihering’s Antwren (*Myrmotherula iheringi*), version 1.0. Birds of the World. https://doi.org/ 10.2173/bow.iheant1.01
